# ALCAM-mediated synapses between DC1 and CD8 T cells are inhibited in advanced lung tumors

**DOI:** 10.1101/2023.10.18.562940

**Authors:** Luciano G. Morosi, Giulia M. Piperno, Sonal Joshi, Roberto Amadio, Simone Vodret, Lucía López-Rodríguez, Federica Benvenuti

## Abstract

Conventional type 1 dendritic cells (DC1) control anti-tumoral CD8 T responses, in lymph nodes and tumor tissues. T-cell activation depends on the establishment of a tight physical interaction with antigen-presenting cell, the immunological synapse (IS). The molecular determinants of DC1-CD8 IS in tumor tissues and how they are regulated during cancer progression remain poorly investigated. Using a reporter for DC1 in a genetic model of non-small cell lung cancer (KP-XCR1^venus^) we show that IS in lung tissues are abundant and productive at early stages of tumor development but progressively diminish in advanced tumors. Transcriptional profiling and flow cytometry of lung resident DC1 identified a module of adhesion molecules downregulated in advanced tumors. We focused on ALCAM and LFA-1, ligands for CD6 and ICAM-1 on T cells, to investigate their role and functional impact. By immobilizing single receptor agonists on artificial cell surfaces, we demonstrate that ALCAM and LFA-1 are sufficient to trigger cytoskeletal remodeling in early tumor DC1, whereas late tumors DC1 are not responsive. Blocking ALCAM-CD6 interactions in functional assays impairs the acquisition of effector functions in CD8 T cells. Together these data highlight that adhesion molecules required to establish IS in early, immune-reactive, tumors are targeted during tumor progression blunting cross-talk within the IS.

## Introduction

The presentation of tumor-derived antigens by antigen-presenting cells is a key factor in priming T cells in lymph nodes and to maintain effector functions and polarization of T cells in tumor tissues. Type 1 dendritic cells (DC1) are uniquely efficient in capturing and cross-presenting cancer antigens to CD8 T cells and their function and regulatory mechanism in lung cancer have been intensely investigated (1–5). For efficient transmission of signal 1 (MHC-peptide complexes), signal 2 (costimulatory molecules) and signal 3 (instructing cytokines), T cell and DC must establish a functional adhesive structure, the IS, to support proper inter-cellular communication (6,7). *Ex vivo* analysis identified complex and dynamic reorganization of the T-IS, encompassing stabilization by adhesion molecules, recruitment of cytoskeletal components and membrane receptors, spatial re-organization of signaling hubs, and polarization of secretory vesicles (8–12). Although less intensely, the DC-IS has been investigated, mostly using surrogate models of dendritic cells differentiated from the blood or bone marrow. These studies identified spatial reorganization of membrane receptors (13), cytoskeletal regulators (14–16), and secretory vesicles (17,18), in addition to changes in the cell surface mechanical properties (19) and epigenomic remodeling (20). Dynamic imaging of DC-T interaction in tissues provided an additional layer, documenting the kinetics of DC-T cells contact that underly programming of effector T cells in lymph nodes (21,22). Further insight into the receptor-ligand pairs regulating DC-T interactions has come from an elegant methodology (LIPSTIC) that unveiled the dynamics of CD40-CD40L interactions in DC2-CD4 IS during immunization (23). Additionally, sequencing of physically interacting immune cells (PIC-seq), in a transplantable lung cancer model identified a similar cluster of CD4 T helper cells associated with antigen-presenting cells as the dominant immune hub controlling anti-tumoral responses (24). In human hepatocellular carcinoma, triads composed of PD-1^+^CXCL13^+^ helper CD4 T cells, mRegDCs and CD8 T cells are critical for the differentiation of tumor-specific progenitors exhausted following checkpoint inhibitors (24,25). Intra-tumoral resident DC1 actively control the formation of these hubs, by attracting CD8 T cell to the stromal tumor regions through CXCL9 and CXCL10 (26–28), thereby promoting expansion of tumor-specific TCF1^+^ stem-like CD8 T cells (28), a mechanism that was recently shown to be impaired by prostaglandin E2 (PGE2) (29). Given the emerging relevance of immune interacting hubs in cancer tissues, understanding the molecular determinants implicated in controlling the stability of DC1-CD8 T cell interactions and their changes during tumor progression becomes an important open question.

Here, we used a model of genetically driven non-small cell lung cancer (NSCLC) recapitulating features of the human disease to focus on resident DC1 of lung tissues to explore their ability to establish productive interaction with CD8 T cells at two stages along tumor development. By combining a reporter mouse line to visualize DC1 in intact lung tissues during tumor progression and imaging of IS formed by primary DC1 isolated from lung tissues, we demonstrate that incipient lung tumors support active IS formation owing to elevated expression of adhesion molecules in DC1. Tumor progression blocks the expression of LFA-1 and ALCAM, reducing the ability to stably interact with CD8 T cells and transmit the signals required for their full differentiation.

## Materials and Methods

### Mice

Mice strains used in this study were maintained in sterile isolators at the International Centre for Genetic Engineering and Biotechnology (ICGEB) animal BioExperimentation facility. Wild type C57BL/6 animals were purchased from ENVIGO Laboratories. XCR1-Venus (*XCR1^Venus/+^*) (30) mice were kindly provided by Dr. Wolfgang Kastenmuller (University of Wuerzburg, Germany). Mice lines B6.129P2-*Trp53^tm1Brn^*/J (p53LoxP, *Trp53^fl/fl^*) and B6.129S4-*Kras^tm4Tyj^*/J (*Kras^LSL-G12D^*) were obtained from The Jackson Laboratory (cod 008462 and 008179, respectively) and initially crossed to obtain the KP inducible mouse line (*Kras^LSL-G12D/+^; Trp53^fl/fl^*). KP-XCR1^Venus^ strain (*Kras^LSL-G12D/+^; Trp53^fl/fl^; XCR1^Venus/+^*) was obtained by crossing the KP and XCR1-Venus colonies. OT-I C57BL/6-Tg(Tcra-V2/Tcrb-V5) animals were from Jackson Laboratories. OT-I-TdTomato mice were a kind gift of Dr. Kastenmuller. The study was approved by ICGEB’s board for animal welfare and authorized by the Italian Ministry of Health (approval number 459/2022-PR, issued on 22/07/2022). Animal care and treatment were conducted with national and international laws and policies (European Economic Community Council Directive 86/609; OJL 358; December 12, 1987). All experiments were performed in accordance with the Federation of European Laboratory Animal Science Association (FELASA) institutional guidelines and the Italian law.

### Induction of lung tumors

We used two mice models of lung adenocarcinoma: autochthonous and transplantable.

In the autochthonous model, 8-10 weeks old KP or KP-XCR1^Venus^ mice were inoculated intratracheally with 2.5 x 10^7^ infectious particles of a replication-deficient adenoviral vector carrying the Cre recombinase gene under the CMV strong promoter (Ad-CMV-iCre, Vector Biolab), in order to induce tumor nodules in the lung. Mice were sacrificed after either 4 or 8 weeks post adenovirus inoculation. In the transplantable model, the KP cell line LG1233 (kindly provided by Dr. Tyler Jacks) was maintained in complete DMEM media (Gibco) as previously described (3), and routinely tested for mycoplasma contamination. In order to induce lung tumor nodules in WT or XCR1-Venus mice, 50,000 KP cells were injected intravenously in 100 µL PBS. Mice were sacrificed after either 8 or 28 days post tumor inoculation. Tumor formation in lung tissues was performed as follows: after sacrifice, healthy and tumor-bearing mice were intracardiac perfused with PBS to remove the vascular component from lungs. Tissues were then harvested and fixed in formaldehyde 10% over-night (ON), and paraffin embedded following standard procedure. 5 µm consecutive sections were dewaxed, rehydrated and stained with the hematoxylin/eosin (H&E; Bio-Optica, Milano Spa). Images were acquired in a Leica DFC450 C microscope.

### Lung immunofluorescence

After sacrifice, healthy and tumor-bearing KP-XCR1^Venus^ lungs were harvested after PBS intracardiac perfusion. Then, PBS 1% paraformaldehyde (PFA) was intratracheally perfused to fix the lung tissue. Left lobe was then placed in PBS 4% PFA ON, and subsequently passed in PBS 15-30% sucrose. Finally, lungs were embedded in Killic (O.C.T., Bio-Optica) and 5 µm sections were obtained in a cryostat. Lung sections were defrosted and immediately fixed with PBS 4% PFA for 10 minutes. Afterwards, cryosections were washed twice with Washing buffer (PBS, 1% BSA, 0.1% Triton X-100) for 5 minutes at RT, and blocked with Blocking buffer (Washing buffer, 5% mouse serum) for 40 minutes at RT. Then, lungs were incubated ON at 4°C with a rat anti-mouse CD8 antibody (4SM15, eBioscience) in Blocking buffer. After ON incubation, sections were washed twice with Washing buffer for 5 minutes at RT, and incubated for 4 hours at RT with a goat anti-rat-AF647 secondary antibody (polyclonal, Invitrogen). After secondary antibody, sections were washed with PBS for 5 minutes at RT, and nuclei were stained with Hoechst 33342 (Invitrogen) in PBS for 20 minutes at RT. Lastly, lung sections were washed twice with PBS for 2 minutes at RT and mounted with Mowiol 40-88 (Sigma-Aldrich). Confocal microscopy images were acquired using a Zeiss LSM 880 instrument, and subsequently analyzed in ImageJ 1.54f software.

### DC1 isolation from lung tissues

Healthy and tumor-bearing lungs were harvested after PBS intracardiac perfusion. Tissues were mechanically cut and digested in complete media containing Collagenase type 2 (265 U/mL; Worthington) and DNase (250 U/mL; Thermo scientific) at 37 °C for 30 minutes. Digestion was then stopped by adding EDTA (10 mM; Invitrogen), tissue pieces were passed through a 18G needle and then filtered using 100 µm cell strainers (Corning) to obtain single-cell suspensions. Red blood cells were eliminated with ACK Lysing buffer (Gibco). Afterwards, CD11c^+^ cells were isolated using magnetic CD11c-microbeads (Miltenyi Biotec) following manufacturer’s instructions. Isolated cells were then stained with anti-mouse antibodies in FACS buffer (PBS, 1% BSA) for 30 minutes at 4°C: CD11c-APC (N418), XCR1-BV650 (ZET), CD172/Sirpα-PE-Dazzle 594 (P84) purchased from BioLegend, and CD170/SiglecF-BB515 (E50-2440) and MHC-II (I-A/I-E)-BV711 (M5/114.15.2) purchased from BD. After washing, stained cells were resuspended in PBS 2% FBS and DC1 were purified in a FACS Aria II Cell Sorter (BD).

### *Ex vivo* DC1-CD8 T-cell conjugates

Sorted lung DC1 were loaded with 10 nM MHC class I-restricted OVA peptide (OVA_257-264_, SIINFEKL) in complete IMDM media (Gibco) for 1.5 hours at 37°C in a Eppendorf tube. Afterwards, they were plated in fibronectin-coated round coverslips for 30 minutes at 37°C. After washing to remove not attached cells, naïve or effector CFSE-labelled OT-I (2 µM CFSE, BioLegend) were added to coverslips in complete RPMI media for 30 minutes at 37°C. After incubation, cells were washed with PBS and fixed in PBS 4% PFA for 5 minutes at RT. Afterwards, cells were permeabilized and blocked in Blocking buffer (PBS, 0.2% BSA, 0.05% saponin, 5% horse serum) for 40 minutes at RT. Then, cells were washed with PBS for 5 minutes at RT and F-actin was stained with Phalloidin-AF555 (Invitrogen) in Blocking buffer for 30 minutes at RT. After washing with PBS for 5 minutes at RT, nuclei were stained with Hoechst 33342 (Invitrogen) in PBS for 20 minutes at RT. Lastly, coverslips were washed with PBS for 2 minutes at RT and mounted with Mowiol 40-88 (Sigma-Aldrich). Confocal microscopy images were acquired using a Zeiss LSM 880 instrument, and subsequently analyzed in ImageJ 1.54f software. Actin distribution in isolated DC1 was calculated using an available macro for ImageJ (31). Naïve OT-I cells were isolated from lymph nodes using magnetic CD8-microbeads (Miltenyi Biotec) following manufacturer’s instructions. Effector OT-I were obtained by co-culturing naïve OT-I for two days with bone marrow-derived DCs (BMDCs), which were previously pulsed for 3 hours with 30 nM SIINFEKL and 1 µg/ml poly(I:C). BMDCs were obtained by differentiating freshly-isolated bone marrow cells in complete IMDM media 30% GM-CSF for 7 days.

### OT-I proliferation assays

Sorted lung DC1 were plated in tissue-treated 96-well plates (5,000 DC1 per well) in complete IMDM for 3 hours at 37°C, in the presence of SIINFEKL at different concentrations (0; 0.3 and 1 nM) and 1 µg/ml poly(I:C). After DC1 were washed, 50,000 naïve CellTrace Violet (CTV)-labelled OT-I cells (5 µM CTV, Invitrogen) were added to each well in complete RPMI media and kept at 37°C for 72 hours. For ALCAM/LFA-1 interfering experiments, recombinant mouse Fc-chimera proteins CD6 (rCD6, R&D Systems) and/or ICAM-1 (rICAM1, R&D Systems) were added to the co-culture at a final concentration of 2 µg/ml. As control, the correspondent recombinant human IgG1 Fc molecule (R&D Systems) was used at the same concentration.

After 48 hours of co-culture, aliquots of conditioned media were collected for measuring IL-2 and IFNγ by ELISA (ELISA MAX, BioLegend). Cells were washed and stained with anti-mouse CD3ε-FITC (145-2C11) and CD8-APC (53-6.7), purchased from BioLegend, in FACS buffer for 30 minutes at 4°C. Afterwards, cells were washed with PBS and stained with LIVE/DEAD Fixable Aqua Dead Cell Stain Kit (Invitrogen) according to manufacturer’s instructions. Finally, cells were fixed in FACS-Fix buffer (PBS, 1% PFA), and CTV dilution was analyzed in a FACS Celesta Cell Analyzer (BD). Data generated were further analyzed in FlowJo software (10.8.1 version, BD).

### Flow cytometry analysis of lungs and lymph nodes

Lungs from tumor-bearing mice (autochthonous and transplantable models) were harvested and processed to obtain single-cell suspensions as previously described. Mediastinal lymph nodes (medLNs) were obtained previous to PBS intracardiac perfusion, and smashed through 100 µm cell strainers using a syringe plunger to obtain single-cell suspensions. Fc receptors were blocked with anti-mouse CD16/32 antibodies (TruStain FcX, BioLegend) in FACS buffer for 10 minutes at 4°C. Single-cell suspensions were then washed and stained with anti-mouse antibodies in FACS buffer for 30 minutes at 4°C, as follows: lungs: CD45-APC/Fire750 (30-F11), CD19-FITC (6D5), CD3ε-FITC (145-2C11), CD45R/B220-FITC (RA3-6B2), NK1.1-FITC (PK136), CD11b-BV421 (M1/70), CD170/SiglecF-PerCP/Cyanine5.5 (E50-2440), CD11c-BV785 (N418), MHC-II (I-A/I-E)-AF700 (M5/114.15.2), XCR1-BV650 (ZET), and CD172/Sirpα-PE-Dazzle 594 (P84), purchased from BioLegend; medLNs: CD19-FITC, CD3ε-FITC, CD45R/B220-FITC, NK1.1-FITC, CD11b-BV421, CD11c-BV785, MHC-II (I-A/I-E)-AF700, and XCR1-BV650. Also, anti-mouse antibodies for adhesion molecules were used, as follows: CD273/PD-L2-PE (TY25), LFA-1 (CD11a/CD18)-APC (H155-78), and CD44-PE (IM7) purchased from BioLegend, and CD166/ALCAM-APC (eBioALC48) purchased from Invitrogen. To quantify lung CD3^+^CD8^+^ absolute numbers, CD45-APC/Fire750, CD3ε-PerCP/Cyanine5.5 (145-2C11) and CD8a-APC (53-6.7) antibodies were used (BioLegend). After staining, cells were washed with PBS, stained with LIVE/DEAD kit as previously described and fixed in FACS-Fix buffer. Cells were analyzed in a FACS Celesta Cell Analyzer (BD). To quantify DC1 and CD8 T cell absolute numbers, CountBrigh Absolute Counting Beads (Invitrogen) were added to samples before being analyzed. Data generated were further analyzed in FlowJo software (10.8.1 version, BD).

### Coating of beads with ALCAM and LFA-1 ligands

Polybead amino 6 µm-diameter microspheres (Polysciences) were prepared to covalently couple rCD6 or rICAM1. Briefly, beads were washed thrice with PBS and activated with glutaraldehyde 8% in PBS for 3 hours at RT, mixing continuously in an orbital rotor. Beads were then washed thrice with PBS to remove glutaraldehyde, and afterwards incubate over-night at 4°C with 2 µg/ml of either rmCD6 or rmICAM1 in PBS, in an orbital rotor.

After washing with PBS to remove any traces of unbound protein, beads were blocked with PBS 1% BSA for 30’ at RT, with continuous mixing in an orbital rotor. Lastly, after washing with PBS beads were resuspend in PBS 1% BSA (50,000 beads/µl) and kept at 4°C until use.

### Lung DC1-beads conjugates

Sorted lung DC1 were plated in fibronectin-coated round coverslips for 30 minutes at 37°C, in complete IMDM media. After washing to remove not attached cells, rCD6- or rICAM1-coated beads were added to coverslips in complete IMDM media for 30 minutes at 37°C (3:1, bead:DC1 ratio). After incubation, cells were washed, permeabilized, blocked, stained for F-actin and nuclei, and finally mounted in Mowiol 40-88, as previously described. Confocal microscopy images were acquired using a Zeiss LSM 880 instrument, and subsequently analyzed in ImageJ 1.54f software.

### ALCAM staining in DC1-CD8 T-cell conjugates

*Ex vivo* DC1-CD8 T-cell conjugates were prepared in fibronectin-coated round coverslips as previously described, using sorted XCR1-Venus early (d8) DC1 and naïve OT-I-TdTomato cells. After incubation, cells were washed once with PBS and stained with anti-mouse CD166/ALCAM-APC (eBioALC48, Invitrogen) antibody in FACS buffer for 30 minutes at 4°C. Afterwards, cells were fixed in PBS 1% PFA for 15 minutes at RT, followed by permeabilization and blocking in Blocking buffer (PBS, 0.2% BSA, 0.05% saponin, 5% horse serum) for 40 minutes at RT. Finally, nuclei were stained with Hoechst 33342 (Invitrogen) in PBS for 20 minutes at RT. Lastly, coverslips were washed with PBS for 2 minutes at RT and mounted with Mowiol 40-88 (Sigma-Aldrich). Confocal microscopy images were acquired using a Zeiss LSM 880 instrument. ALCAM integrated density at the synapse or at the opposed side in XCR1-Venus DC1 was calculated in ImageJ 1.54f software, an normalized by the total ALCAM density of the cell.

### Datasets analysis

Microarray data from purified control and KP-lung DC1 was previously generated in our laboratory (3), is publicly available in Gene Expression Omnibus (GEO) under accession code GSE119574. scRNA-seq data from control and KP lungs were obtained from GEO under accession code GSE131957 (2). Briefly, we downloaded from the GitHub repository associated to the publication (https://github.com/effiken/Maier_et_al_nature_2020) the R object file containing expression values and cell annotations. Cell adhesion molecules’ genes were obtained from the Kyoto Encyclopedia of Genes and Genomes (KEGG) Pathway Database, especially focalizing in those adhesion molecules involved in immune system cell-to-cell interactions (https://www.genome.jp/pathway/hsa04514). Genes were confronted with our microarray data to find those differentially expressed (DEGs) between control and KP DC1 (p-value<0.05). Afterwards, DEGs were validated in the scRNA-seq dataset. Normalized expression values for selected adhesion molecules genes (*Alcam*, *Itgb2*, *Itgal*, *Cd44*) in steady-state mouse dendritic cells were retrieved from the Immunological Genome Project (ImmGen; https://www.immgen.org/) Microarray V1 Dataset. Expression values were log2 transformed and z-score normalized for visualization. An “Adhesion score” for each dendritic cell population was calculated as the mean z-score between these adhesion molecules genes.

### Statistical analysis

GraphPad Prism version 8 (GraphPad Software Inc.) was used for statistical analysis. The number of replicates for each experiment is presented in the respective figure legends. Two groups were compared with two-tailed Student’s t-test for paired or unpaired data. For multiple comparisons, one-way or two-way ANOVA followed by Tukey’s or Dunnett’s post-tests (parametric analysis), or Kruskal-Wallis followed by Dunn’s post-test (nonparametric analysis) were used. P values lesser than 0.05 were considered significant.

## Results

### Visualization of DC1-T immune synapses along progression of autochthonous tumors

To investigate the morphology and distribution of DC1-CD8 T cell contacts in the context of progressive lung tumors, we generated a variant of the KP genetic model of NSCLC (*Kras^LSL-G12D/+^*; *Trp53^fl/fl^*) crossed to the XCR1^Venus/+^ reporter (30), to unequivocally visualize DC1 in tissues (KP-XCR1^venus^) (**Fig. 1A**). Lung tumors were induced by intratracheal administration of Cre recombinase and tissues were harvested after 4 weeks (corresponding to early adenomas) and 8 weeks (corresponding to established adenocarcinoma) (**Fig. S1A**). Lung tissues of control, not induced mice, showed few sparse DC1 distributed evenly in the lung parenchyma. At early time points after tumor induction, we noted a significant increase in DC1 numbers, which appeared aggregated in small patches (**Fig. 1B** and **C**). At late time points, the number of DC1 had diminished and cells were found either sparse or clustering at the border of tumor nodules (**Fig. 1B** and **C**). The increment in DC1 at 4 weeks was paralleled by an increase in the numbers of CD8 T cells, while their numbers dropped at 8 weeks post-tumor induction (**Fig. 1D**). We then quantified the number and multiplicity of IS in the lungs and the maximum angle of Venus fluorescence at the cell-cell interface, as an estimate of the interaction’s tightness. Healthy lungs displayed very few DC-T monogamous conjugates (i.e., featuring 1 DC and 1 T cell) (**Fig. 1E-G**). Remarkably, a higher number of conjugates and a larger fraction of DC1 were found to be engaged with CD8 T cells in tissues carrying early tumor lesions (**Fig. 1E** and **F**). Moreover, a high fraction of DC1 were in contact with multiple CD8 T cells (**Fig. 1G**) and the interactions were tighter, as measured by the Venus angle at the synapse (**Fig. 1H**). In contrast, established tumors showed fewer conjugates and most interactions were monogamous and less tight (**Fig. 1E-H**).

**Figure 1.**
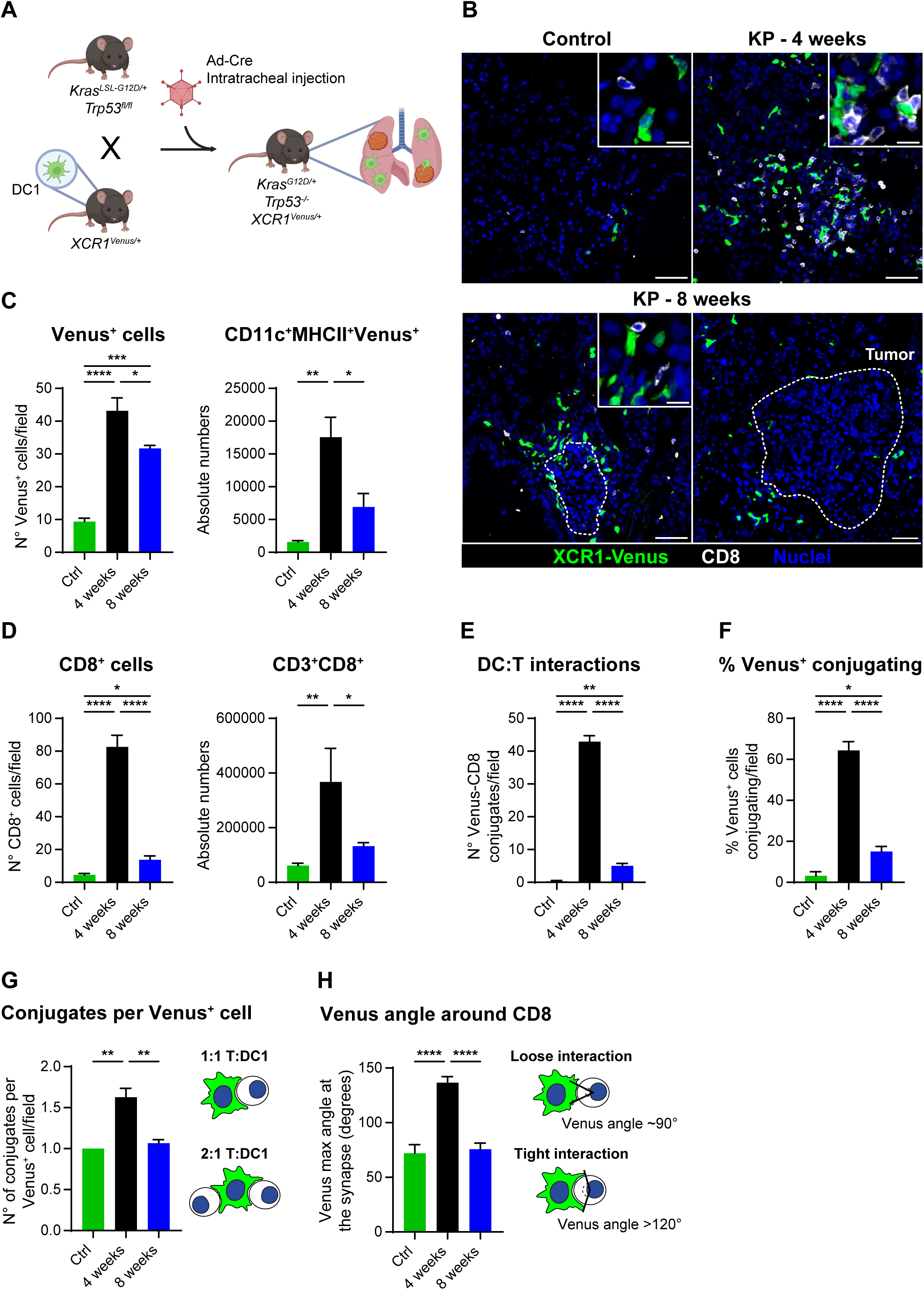
Defective DC1-CD8 T cell interactions in advanced KP lung tumors. **A.** KP mice strain (*Kras^LSL-G12D/+^*; *Trp53^fl/fl^*) was crossed with the XCR1-Venus reporter (*XCR1^Venus/+^*) to generate the KP-XCR1^Venus^ line. Tumors were induced by intratracheal Cre-Recombinase Adenovirus (Ad-Cre) administration (created with BioRender.com). **B.** Representative confocal planes showing DC1-CD8 T-cell conjugates in 5µm lung cryosections from healthy (control) or tumor-bearing lungs (KP) after 4- or 8-weeks post Ad-Cre administration. Lungs were stained for CD8 T cells (white) and nuclei (blue), while XCR1^+^ DC1 expressed the Venus reporter (green). Scale bar large images, 50 µm; scale bar insets, 6 µm. **C.** Number of Venus^+^ DC1 quantified in tissue sections (left) and by flow cytometry (right). **D.** Number of CD8^+^ cells quantified in tissue sections (left) and by flow cytometry (right). **E-F.** Number of total DC-T cell interactions (**E**) and percentage of Venus^+^ DC1 engaged in IS (**E**). **G.** Multiplicity of interactions for each Venus^+^ cell found in the lung tissue. **H.** Tightness of the DC1-CD8 T cell interactions, measured as the Venus maximum angle at the synapse (around a CD8 T cell). Data represent mean ± SEM, one representative out of two independent experiments is shown. For Venus^+^ DC1 (**C**) and CD8^+^ (**D**) absolute numbers n = 3 mice per group, Kruskal-Wallis followed by Dunn’s post-test. For cryosections analysis in **C-G**, n = 6-8 planes were analyzed, pooled from 3 left lobes acquired per group. In **H**, for Venus angle quantification, n = 7 in Control, n = 42 in 4 weeks, and n = 28 in 8 weeks DC1-CD8 interactions were analyzed. One-way ANOVA followed by Tukey’s post-test. *p<0.05 **p<0.01 ***p<0.001 ****p<0.0001.

In summary, these observations in the KP-XCR1^Venus^ reporter unveil that tissue-resident DC1 actively scan the CD8 T cells repertoire at early stages of tumor development in autochthonous lung tumors. However, this capacity is transient and IS are rarely observed in lung tissues carrying established tumors.

### Lung tumors impair the formation of productive DC-T cell synapses *ex vivo*

To dissociate tissue factors that may impede DC-T cell encounter from DC1 intrinsic changes impairing the IS, we next isolated DC1 from lung tissues to establish synapses *ex vivo*. To facilitate the recovery of sufficient numbers of primary cells from tissues and to precisely control the tumor stage, we grafted KP tumors orthotopically by intravenous injection, a model that recapitulates progressive dysfunction in the DC1 compartment (3). Tissue were harvested after 8 or 28 days, corresponding to early dysplasia, or established adenocarcinoma (**Fig. S1A**). Cell-sorted DC1 were pre-pulsed with OVA class-I peptide to bypass defective antigen uptake and processing and mixed with functional, OVA-specific CD8 T cells (OT-I) (**Fig. 2A**). We used both naïve and effector OT-I, to mimic priming in lymph nodes and secondary interactions in tissue, respectively. DC1 isolated from early tumors induced significantly more conjugates with naïve OT-I than those of control tissues, indicating enhanced adhesive properties triggered by early tumors, intrinsically in DC1 (**Fig. 2B** and **C**). In contrast, DC1 isolated from advanced tumors were strongly impaired in forming IS. Furthermore, the surface of interaction (measured as the actin maximum angle at the synapse) and the actin thickness at the synaptic interface were significantly diminished in late tumor DC1, as compared to control and early tumor DC1, suggesting defective signaling and cytoskeletal remodeling (**Fig. 2B** and **C**). In the case of effector OT-I, both control and early DC1 formed comparable numbers of conjugates, nevertheless, this was markedly blunted in late DC1 (**Fig. S1B**), suggesting that the latter are impeded also to interact with activated CD8 T cells. Given the changes observed in the IS actin distribution between late and control/early DC1, we examined the structure of actin at basal level. Even though the actin distribution across the cell was comparable between control and tumor-conditioned DC1 (**Fig. S1C**), late DC1 presented a basal ticker cortical actin than control/early DC1 (**Fig. S1D**). This suggests that DC1 in late tumors may have a different stiffness, impacting on the ability to engage TCR signaling.

**Figure 2.**
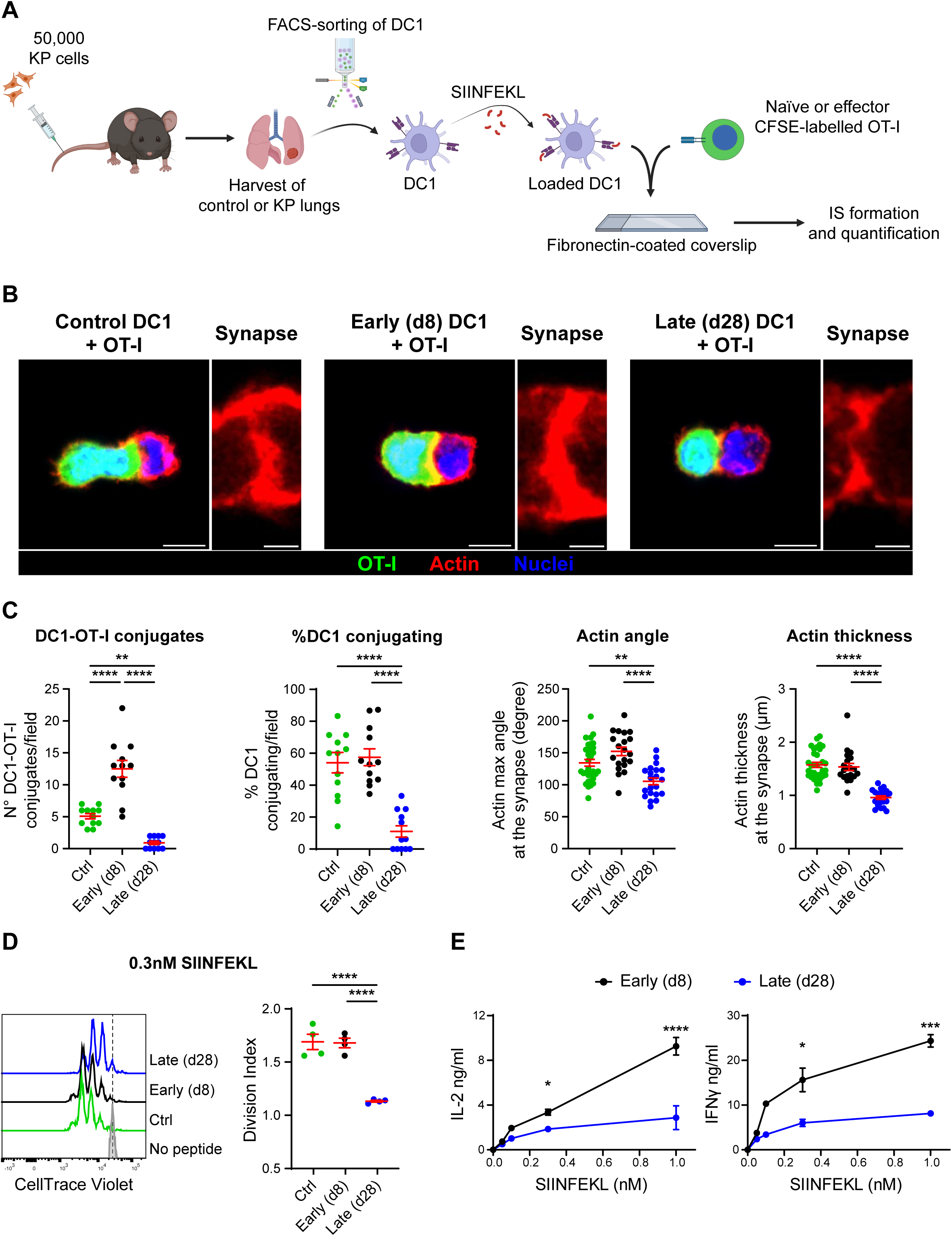
Tumor-conditioned lung DC1 are impaired to establish productive IS with CD8 T cells. **A.** Wild-type male mice were injected intravenously with 50,000 KP cells to generate tumor nodules in the lung. DC1 were isolated from control or tumor-bearing tissues (at day 8, early, or 28, late) by fluorescence-activated cell sorting, then they were loaded *ex vivo* with class-I restricted OVA-peptide (SIINFEKL) and mixed with CFSE-labelled naïve or effector OT-I cells on fibronectin-coated coverslips (created with BioRender.com). **B.** Representative confocal planes showing DC1-OT-I conjugates (scale bar, 6 µm) and the actin structure of the immunological synapse (scale bar, 2 µm). **C.** Quantification of the numbers of DC1-OT-I conjugates generated and percentage of DC1 conjugating (n = 12 planes per group), surface of interaction at the IS (actin maximum angle), and actin thickness at the IS are presented (n = 20-32 DC1-OT-I conjugates per group). One-way ANOVA followed by Tukey’s post-test. **D-E.** Isolated lung DC1 were loaded *ex vivo* with different concentrations of SIINFEKL and mixed with CTV-labelled OT-I cells (1:10 DC1:T cell ratio). **D.** Representative histogram of CTV dilution of OT-I cells (left) and division indexes for each group (right, n = 4 replicates pooled from two independent experiments) after 72 hs of co-incubation (0.3 nM SIINFEKL). One-way ANOVA followed by Tukey’s post-test. **E.** IL-2 (left) and IFNγ (right) produced by OT-I cells after 48 hs of co-incubation, measured by ELISA in the supernatants (n = 3 replicates per SIINFEKL concentration, one representative out of two independent experiments). Two-way ANOVA followed by Tukey’s post-test. Data represent mean ± SEM, one representative out of three independent experiments is shown. *p<0.05 **p<0.01 ***p<0.001 ****p<0.0001.

To correlate these observations to the functional outcome of the interaction, we loaded DC1 with different concentrations of OVA class-I peptide to stimulate CTV-labelled naïve OT-I cells. OT-I cells stimulated by late tumors DC1 proliferated significantly less than those incubated with control or early tumors DC1 (**Fig. 2D** and **Fig. S1E**). Moreover, CD8 T cells primed by late DC1 produced less IL-2 and IFNγ than those primed by early tumors DC1 (**Fig. 2E**). Taken together, our results identify two-stages along tumor progression, the first actively promoting contact formation by DC1 and the second inhibiting them, corresponding to opposite functional outcomes.

### Lung tumors provoke a down-regulation of adhesion molecules in lung DC1

We previously generated a dataset comparing lung resident DC1 at the steady-state to those isolated from advanced lung tumor tissues (3). Focusing on co-stimulatory/co-inhibitory and adhesion molecules that may control interaction with T cells, we identified a set of 46 significantly (p<0.05) expressed genes (20 upregulated, 26 downregulated) (**Fig. 3A**). Co-inhibitory molecules PD-L1 and PD-L2 were among the most upregulated genes, whereas the adhesion molecules CD43, ALCAM (CD166), both subunits of LFA-1 (CD18/CD11a), and CD44 were downregulated in late tumor DC1 (**Fig. 3A**). In a publicly available single-cell RNA dataset of KP tumors (2), we confirmed down downregulation of CD44 and ALCAM paralleled by upregulation of PD-L1 and PD-L2 (**Fig. 3B**). Interestingly, these adhesion molecules are selectively highly expressed in lung resident DC1, according to the ImmGen Database (**Fig. S2A**). We then sought to validate these data by analyzing the expression of selected molecules (ALCAM, CD44, LFA-1, and PD-L2) by flow cytometry on lung DC1. We found a significant loss in the expression of ALCAM, LFA-1 and CD44 in DC1 at late tumor stages, in both the autochthonous and transplantable KP models (**Fig. 3C**). Furthermore, in late DC1, we found a substantial increment in the population of PD-L2^+^ cells (**Fig. 3C**). We examined migratory DC1 (CD11c^+^MHC-II^high^XCR1^+^) in mediastinal lymph nodes (medLNs) of control, early and late tumors to explore whether these changes extended beyond tumour tissues. ALCAM and PD-L2 were reduced and increased, respectively, mirroring what happens in lung DC1, in both endogenous and transplantable tumor models (**Fig. S2B**). CD44 was slightly diminished in the endogenous but not in the transplantable model, whereas LFA was unchanged in the endogenous and reduced in the transplantable model (**Fig. S2B**). These data show that defective IS formation by late tumor DC1 correlates to diminished expression of ALCAM, CD44 and LFA-1, suggesting a direct causal link.

**Figure 3.**
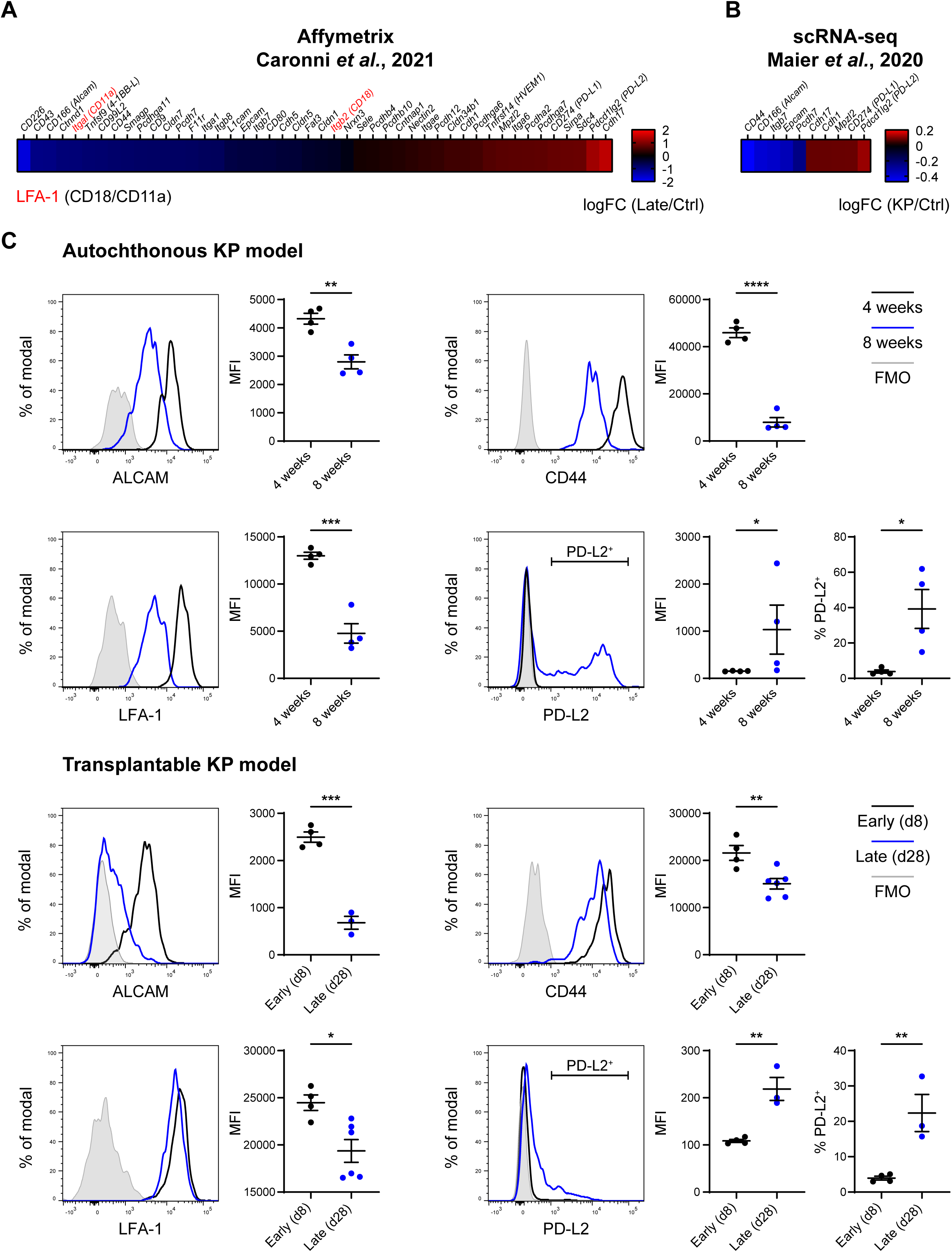
Prominent downregulation of ALCAM and LFA-1 in lung DC1 associated to advanced KP tumors. **A.** Heatmap showing adhesion molecules differentially expressed (p<0.05) in late KP-bearing lung DC1 *vs* control DC1 in our data set (Caronni *et al.*, 2021). **B.** Adhesion molecules significantly modulated in lung late DC1, based on the scRNA-seq dataset from (Maier *et al.*, 2020). **C.** Tumor-bearing lungs were harvested and processed for flow cytometry. In the autochthonous model, lungs were harvested and analyzed after 4- (black line) or 8-weeks (blue line) post Ad-Cre administration (n = 4 mice per group, one representative out of two independent experiments). In the transplantable model, lungs were analyzed at day 8 (black line, n = 4 mice) or 28 (blue line, n = 3-6 mice) after intravenous KP tumors inoculation (one representative out of three independent experiments). Fluorescence minus one (FMO) control staining for each marker is shown in grey. Quantification of the Median Fluorescence Intensity (MFI) or percentage of positive cells in lung DC1 for the indicated markers is presented. Data represent mean ± SEM. Unpaired Student t-test. *p<0.05 **p<0.01 ***p<0.001 ****p<0.0001.

### Engaging ALCAM or LFA-1 induces cytoskeletal rearrangement similar to IS in early DC1

The integrin LFA-1 is a key regulator of synapse formation and TCR signaling in T cells. Interaction with its ligand, ICAM-1 expressed on dendritic cells, triggers inside-out signaling to achieve full TCR signal transduction, T cell activation and degranulation of cytotoxic T cells against their target (32,33). The reciprocal ligand pair, i.e. LFA-1 on dendritic cells engaging ICAM-1 on T cells, was previously reported to promote T cell activation by dendritic cells (34). On the same line, the binding of ALCAM on dendritic cells to CD6 on CD4 T cells has been shown to regulate CD4 T cells activation (35). To directly investigate the role of adhesion molecules in IS formation with CD8 T cells we isolated DC1 from XCR1-Venus mice carrying early transplantable KP tumor and loaded them with peptide before mixing to OT-I-TdTomato, followed by fixation for confocal analysis. We used antibodies to ALCAM as strong expression of LFA-1 in T cells would mask DC1 specific events (36). As shown in **Fig. 4A**, ALCAM is evenly distributed on the surface of DC1, but when these cells form conjugates with OT-I cells the receptor preferentially accumulates at the synaptic side of the cell membrane (**Fig. 4A**). Next, we developed a reductionist approach to selectively trigger LFA-1 and ALCAM on primary lung DC1. To this goal, we prepared 6 µm-polystyrene beads functionalized with recombinant CD6 (rCD6) as ligand for ALCAM, and rICAM1 as ligand for LFA-1. Beads were added to DC1 isolated from early or late tumor tissues and incubated for 30 minutes before washing and processing for confocal analysis (**Fig. 4B**). Control uncoated beads were found in close apposition to the DC1 surface, without triggering membrane remodeling (**Fig. 4C** and **D**). Interestingly, DC1 isolated from early tumors responded to beads carrying rCD6 or rICAM1 by membrane deformation and recruitment around the interacting beads, as measured by the maximum actin angle at the DC1-bead interface (**Fig. 4C** and **D**). Consistent with the loss of expression of the ligand, and similarly to what was observed with whole CD8 T cells, DC1 from late tumor-bearing lungs formed significantly fewer contacts with rCD6- and rICAM1-beads. Moreover, the DC1-bead surface of interaction was smaller and flatter, resembling that formed by control uncoated beads (**Fig. 4C** and **D**). We conclude that engaging single adhesion molecules such as ALCAM or LFA-1 on DC1 from early tumors is sufficient to generate a cellular response that stabilizes the interaction, suggesting their involvement in initial T cell scanning. Consequently, their loss in late tumors may explain, at least in part, defective IS formation.

**Figure 4.**
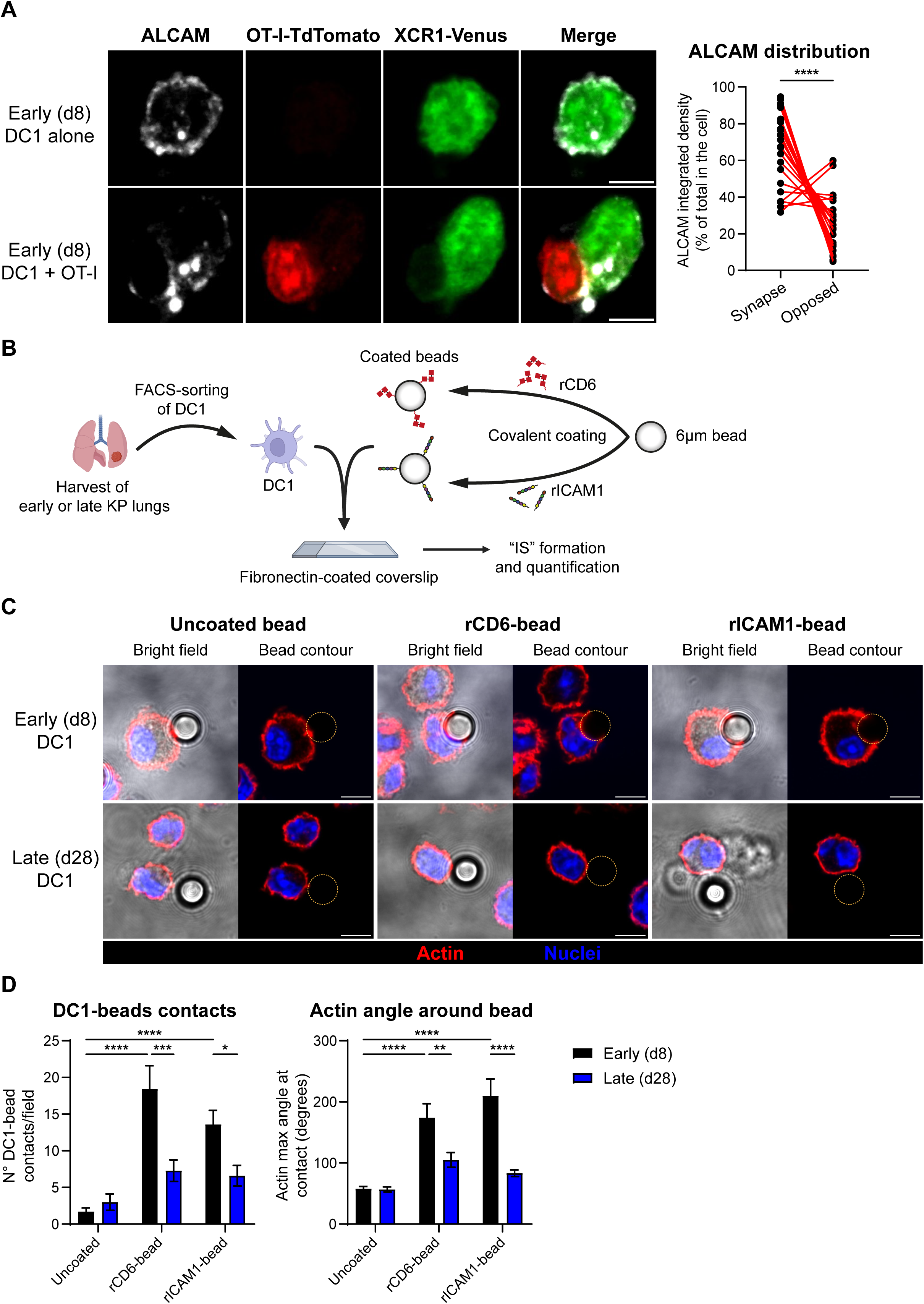
Engaging ALCAM or LFA-1 induce cytoskeletal rearrangement in early DC1. **A.** XCR1-Venus DC1 (green) isolated from early (day 8) tumor-bearing lungs were loaded *ex vivo* with SIINFEKL and mixed with naïve OT-I-TdTomato cells (red) on fibronectin-coated coverslips, and subsequently stained for ALCAM (white). Representative confocal planes showing ALCAM distribution in early DC1 alone (top images) or in DC1-OT-I conjugates (bottom images) (scale bar, 4 µm). Quantification of ALCAM distribution in conjugating DC1, either at the synapse or at the opposed side (n = 24 DC1-OT-I conjugates). Data represent mean ± SEM. Paired Student t-test. **B.** Functionalized beads were generated to study individual molecules involved in IS formation. 6 µm-beads were coated either with recombinant mouse ligands for ALCAM (rCD6), for LFA-1 (rICAM1), or were left uncoated. The beads were incubated with isolated DC1 from early or late KP lungs on fibronectin-coated coverslips (created with BioRender.com). **C.** Representative confocal planes showing DC1-bead conjugates (scale bar, 6 µm). **D.** Quantification of DC1-bead contacts (n = 10 fields per group) and DC1-bead surface of interaction (actin maximum angle, n = 10 DC1-bead conjugates). Two-way ANOVA followed by Tukey’s post-test. Data represent mean ± SEM, one representative out of three independent experiments is shown. *p<0.05 **p<0.01 ***p<0.001 ****p<0.0001.

### Interfering with ALCAM-CD6 interactions in early DC1 prevents T cell activation

Finally, we aimed to directly address the functional significance of ALCAM and LFA-1-mediated interactions on CD8 T cell priming. Thus, we performed T cell activation assays incubating peptide-loaded DC1 from early tumors (the condition that maximally activates CD8 T cells, **Fig. 2D** and **E**) with CTV-labelled naïve OT-I cells. To interfere with the ALCAM-CD6 and LFA-1-ICAM axis, we added to the co-culture soluble rCD6 or rICAM1, individually or in combination, to block the interaction with the respective T cell receptor (**Fig. 5A**). Given that both recombinant proteins are fused with a human IgG1 constant fragment (Fc) to increase their stability, we used the corresponding recombinant Fc (rFc) as a control. In spite of no effect on CD8 T cell proliferation, interfering with ALCAM or LFA-1 (**Fig. 5B**) significantly reduced IL-2 released in the coculture supernatant was found (**Fig. 5C**). In agreement, the effector cytokine IFN-γ was reduced by ALCAM-CD6 interference (**Fig. 5D**).

**Figure 5.**
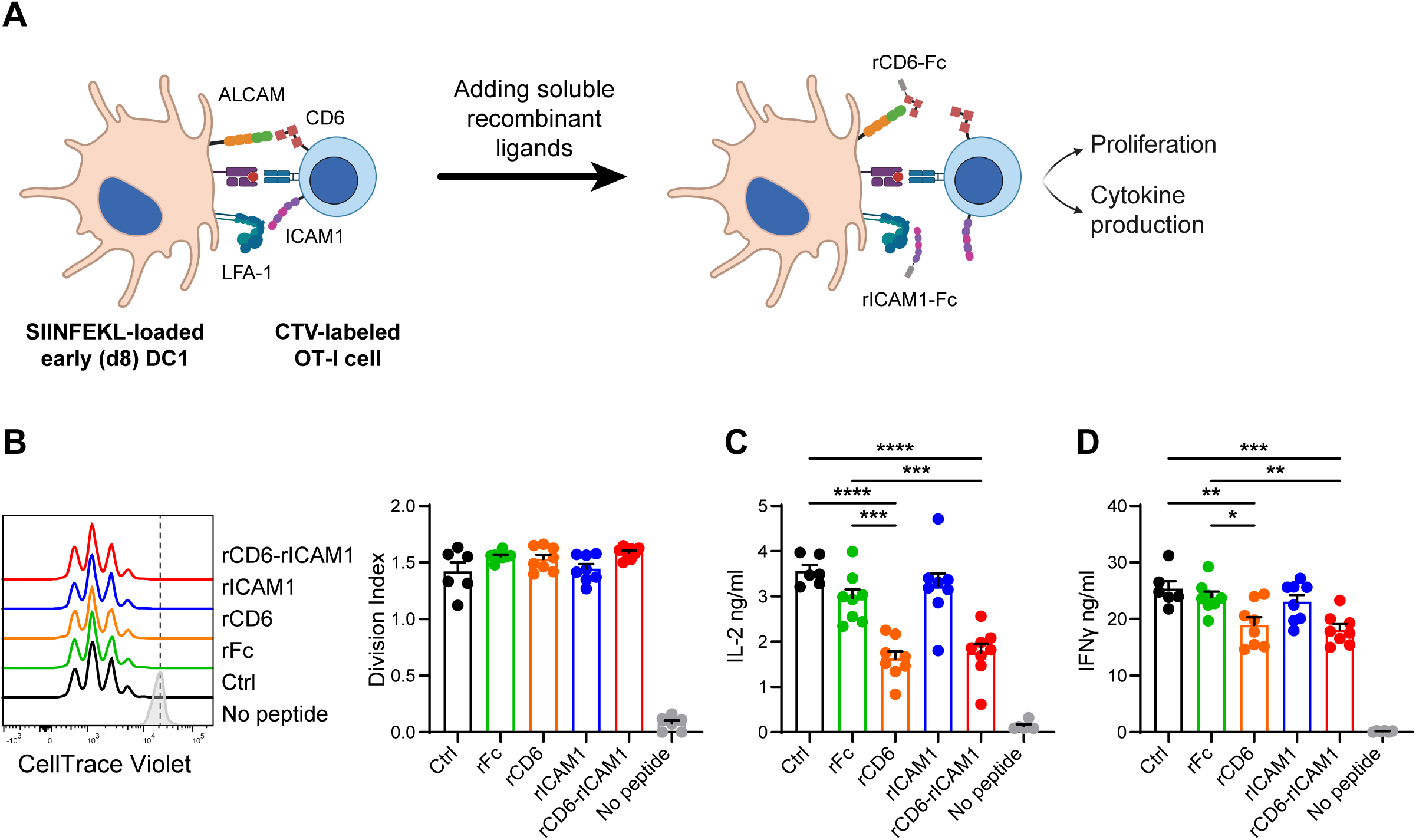
ALCAM expression in lung DC1 is critical to support activation of naïve CD8 T cells. **A.** Isolated lung early (d8) DC1 were loaded *ex vivo* with SIINFEKL and mixed with CTV-labelled OT-I cells (1:10 DC1:T cell ratio) in the presence of soluble recombinant mouse ligands for ALCAM (rCD6) and/or LFA-1 (rICAM1) (created with BioRender.com). **B.** Representative histograms of CTV dilution of OT-I cells (left) and division indexes for each group (right) after 72 hs of co-incubation. IL-2 (**C**) and IFNγ (**D**) produced by OT-I cells after 48 hs of co-incubation, measured by ELISA in the supernatants (n = 6-8 replicates pooled from three independent experiments). Data represent mean ± SEM. Two-way ANOVA followed by Tukey’s post-test. *p<0.05 **p<0.01 ***p<0.001 ****p<0.0001.

These results collectively suggest that blocking ALCAM-mediated interactions does not prevent CD8 T cells from entering into cycling. However, ALCAM-CD6 binding is required to stabilize the interaction and to promote effector functions beyond expansion.

## Discussion

The physical interaction between CD8 T cells and cross-presenting DC1 is a central step in the cancer immune cycle. In addition to interactions occurring in lymph nodes for priming naïve CD8 T cells against tumor-derived neoantigens, secondary interactions in tumor tissues are critical to regulate anti-tumoral immune responses. By focusing on lung tissue-resident DC1, we show that early steps of *in situ* carcinogenesis trigger an active scanning behavior which results in multiple contacts with CD8 T cells, whereas tissues harboring advanced tumors inhibit intercellular DC-T communication. We have identified a module of lung-specific DC1 adhesion molecules involved in facilitating IS formation and signal transmission that is selectively downmodulated in late tumors, suggesting a previously unappreciated mechanism of immune evasion targeting IS stability in tumor tissues.

This study is unique in combining the analysis of DC1-CD8 contacts in tissues, to confocal imaging at higher resolution of IS formed *ex vivo* using DC1 isolated from lung tissues. To facilitate the detection of DC1-IS we crossed a DC1 reporter line (30) to the inducible genetic model of NSCLC, which has proven to faithfully recapitulate the features of the human disease (5,37). This approach has two major advantages. First, it allows precise identification of DC1, overcoming the limitations of labeling promiscuous markers (XCR1, CD103, CD11c) that may be modulated during DCs activation. Second, the tumorigenic process occurs *in situ*, recapitulating the correct dynamics of tumor evolution. By quantifying DC1-CD8 in tissues, we observed a significant increase in clustering at initial tumor stages, corresponding to early adenomas (38), as compared to healthy lungs. We speculate that sterile inflammation triggered by initial tissue damage and recognition of danger-associated molecular patterns, including cGAS-STING activation, may promote pathways supporting cell-cell encounter and interaction. Since the KP-XCR1^Venus^ model harbours very few immunogenic antigens (Research Square [https://doi.org/10.21203/rs.3.rs-2752957/v1]), the contacts captured in fixed tissues may represent events of active scanning, preceding recognition of cognate antigens and IS stabilization. Alternatively, low-affinity neoantigens may be sufficient to engage cells into stable IS, at tumor inception. In both cases, our results unveil that early tumors actively promote surveillance of the tissues by inducing scanning in search for cognate antigens. In contrast, tissues invaded by adenocarcinomas counteract IS formation and/or maintenance. To understand whether tissue or DC1-intrinsic factors were primarily responsible for the changes in the capacity to interact, we isolated DC1 from tumors of different grades to probe the ability to form IS *ex vivo*. Our results strongly suggest DC1 cell-autonomous factors contribute to both enhanced clustering in early stages and failure to stabilize the IS at late stages (**Fig. 6**).

**Figure 6.**
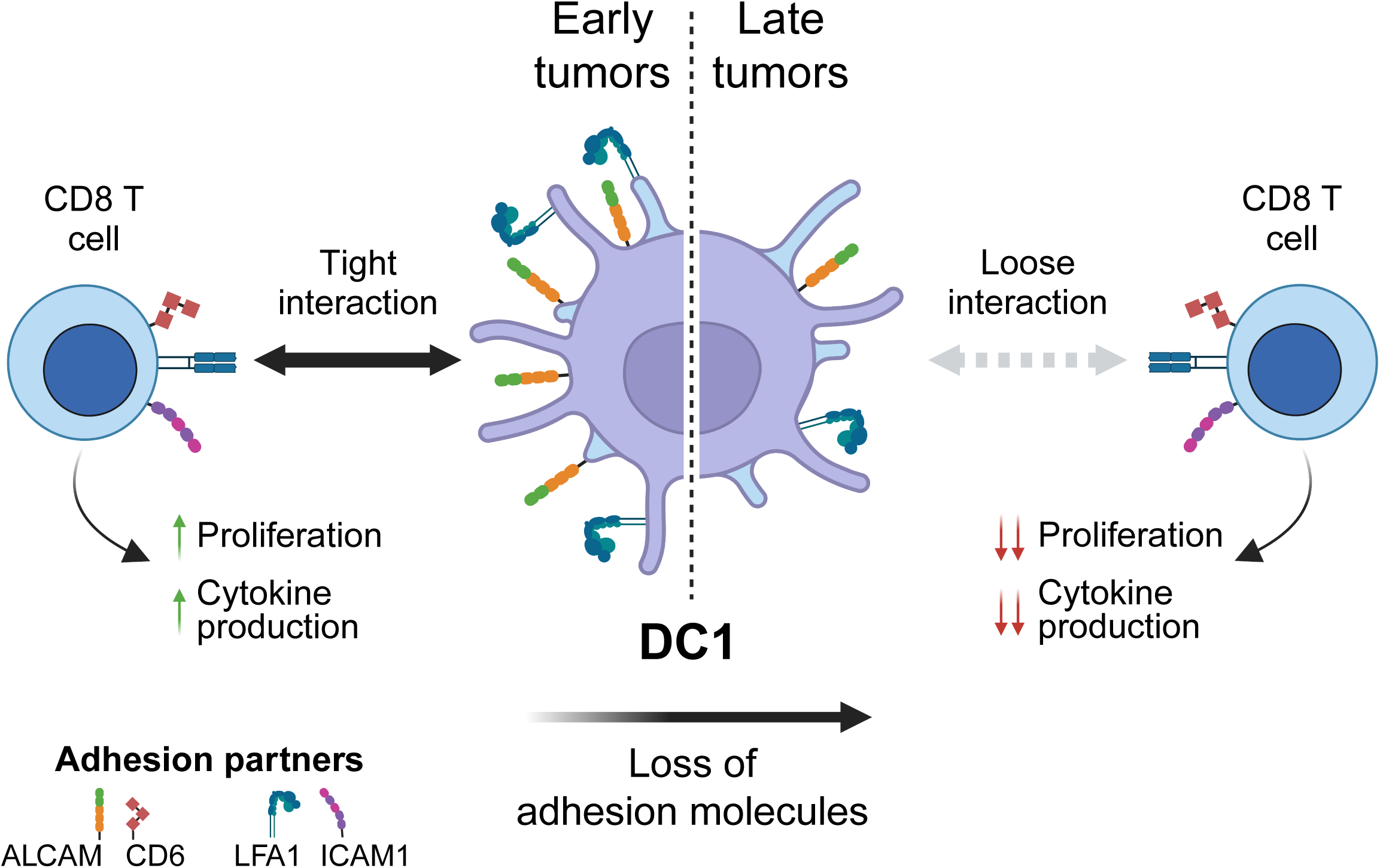
Model of impairment of anti-tumor CD8 responses in lung due to the loss of adhesion molecules in DC1. At early stages of lung cancer, DC1 express high levels of adhesion molecules (like ALCAM and LFA-1, but not only), which allows them to establish productive IS with CD8 T cells to cross-present tumor derived antigens. Nevertheless, along with tumor progression DC1 loss the expression of many of these adhesion molecules, which renders in loose interactions with CD8 T cells, thus impairing the activation of anti-tumor responses (created with BioRender.com).

Activation of tumor-specific CD8 T cells by DC1 is a complex, multistep process including antigen capture and cross-presentation, innate sensing to upregulate costimulatory signals, migration to specific niches to convene with CD8 T cells and, finally, engagement in tight physical interactions. Whether the ultimate step of DC1-CD8 intercellular communication may be modulated, across tumor development and by what mechanism, has been little investigated. Both the initial steps of DC-T scanning prior to TCR-pMHC recognition and those following TCR engagement are largely controlled by adhesion molecules that facilitate membrane proximity and regulate downstream cytoskeletal remodeling to support intracellular signals (39). The role of LFA-1 in T cells is well-documented (40,41), whereas the significance of LFA-1 on DCs is poorly understood and limited to the analysis of DCs generated from bone marrow precursors (34). Using artificial beads reconstituted with single ligands we could isolate the effect of LFA-1 on DC1, demonstrating that it triggers a response involving cellular deformation around the target, accompanied by actin remodeling. However, we cannot definitely conclude on the impact of LFA-1 loss in late tumors DC1, since its blockade has no functional consequence, at least in our assay, suggesting a redundant role. In parallel, we identified ALCAM as highly expressed on control and early tumors lung DC1 and significantly downmodulated in late tumors, both locally and in tumor draining lymph nodes (**Fig. 6**). Beads coated with the ALCAM ligand CD6 are sufficient to trigger conjugation with DC1 and its blockade inhibits effector functions in interacting CD8 cells. Therefore, we propose a non-redundant role for ALCAM in controlling tissue surveillance, by promoting DC-T scanning in lung tissues. Consistently, previous studies demonstrated the importance of the ALCAM-CD6 axis to stabilize the immunological synapse (42) and to promote T-cell priming by human monoDCs (35,43,44). Mechanistically, ALCAM was reported to form a link between CD6 and the actin cortex to strengthen cell adhesion (42,44), consistent with our observation on actin remodeling. Furthermore, blocking ALCAM with an antibody reduces DC migration both *in vitro* and *in vivo*, as well as CD4 T-cell activation *in vitro* by BM-DCs (45). In summary, this study presents valuable tools for studying DC1 in lung cancer tissues, providing initial insights into the mechanism that regulates synapse formation and its suppression during tumor progression.

## Acknowledgements

This work was supported by AIRC IG 21636 to FB. LGM, SJ and LL-were supported by ICGEB Arturo Falaschi pre- and post-doctoral fellowships. RA is supported by Italian Telethon.

## Author contributions

LGM and FB conceived the study, designed experiments, and wrote the manuscript. FB supervised the study. LGM performed experiments, data analysis, and prepared the figures. GMP, SJ, RA and LL-R assisted in the execution of experiments and critical discussion of the data. LGM and RA analyzed gene expression data. SV provided assistance with the management and manipulation of the animal models. All authors read and approved the final version of the manuscript.

