## Supplementary Figures 1 and 2 for "ALCAM-mediated synapses between DC1 and CD8 T cells are inhibited in advanced lung tumors"

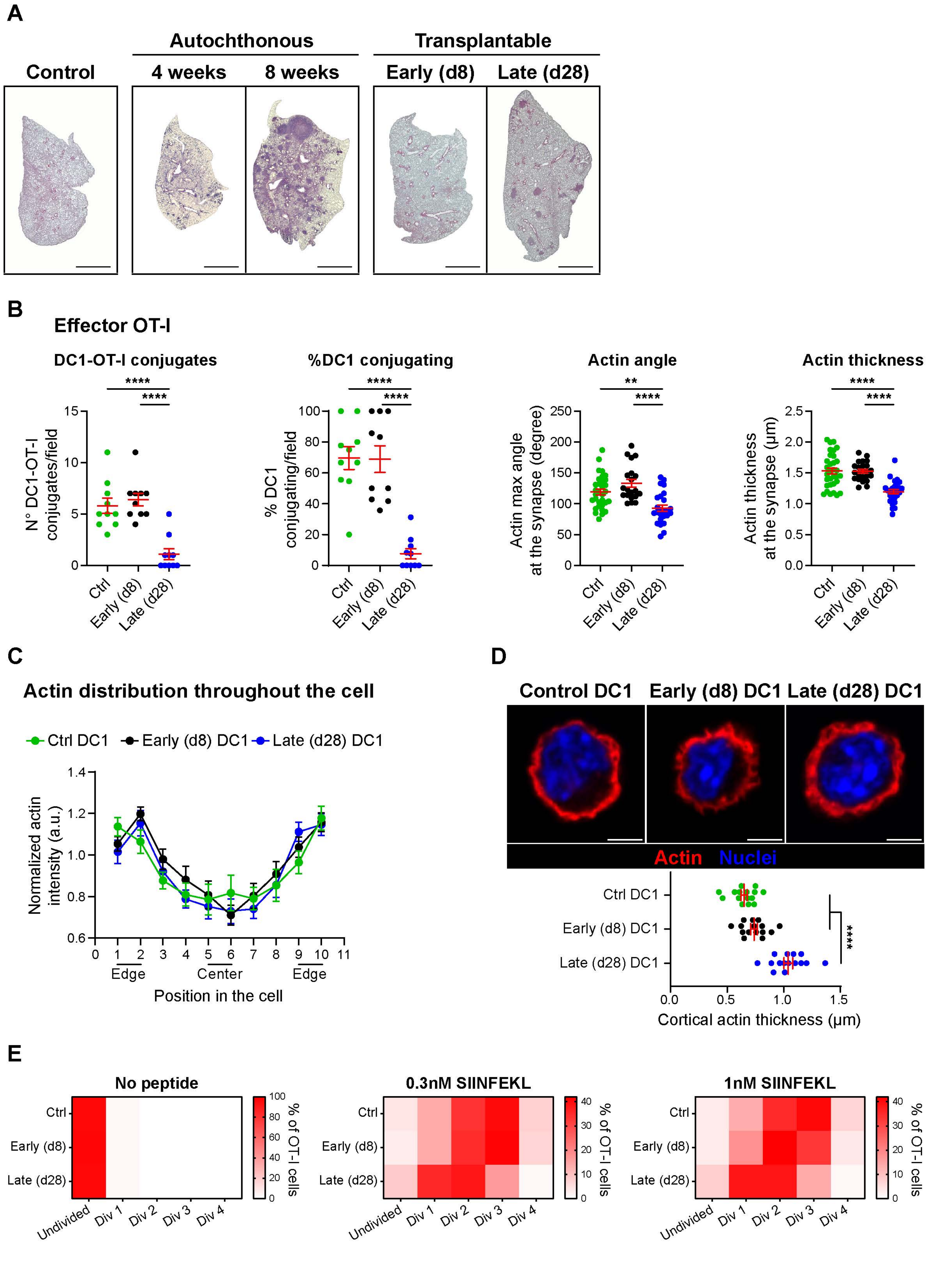
A
Autochthonous
Transplantable
Control
4 weeks
8 weeks
Early (d8)
Late (d28)
B
Effector OT-I
DC1-OT-I conjugates
%DC1 conjugating
Actin angle
Actin thickness
N° DC1-OT-I conjugates/field
% DC1 conjugating/field
Actin max angle at the synapse (degree)
Actin thickness at the synapse (μm)
Ctrl
Early (d8)
Late (d28)
C
Actin distribution throughout the cell
Ctrl DC1
Early (d8) DC1
Late (d28) DC1
Normalized actin intensity (a.u.)
Edge
Center
Edge
Position in the cell
D
Control DC1
Early (d8) DC1
Late (d28) DC1
Actin Nuclei
Cortical actin thickness (μm)
E
No peptide
0.3nM SIINFEKL
1nM SIINFEKL
Undivided
Div 1
Div 2
Div 3
Div 4
% of OT-I cells

Figure S1

**Figure S1. Late DC1 fail to physically interact with CD8 T cells. A.** Representative H&E sections of healthy (control), or tumor-bearing lungs in both KP models used. For the autochthonous model, lungs were harvested after 4- or 8-weeks post Ad-Cre administration; whereas for the transplantable model, after 8 (early) or 28 (late) days post intravenous KP cell line administration (scale bar, 5 mm). **B.** DC1 were isolated from control or tumor-bearing lung tissues, loaded *ex vivo* with SIINFEKL and mixed with CFSE-labelled effector OT-I cells on fibronectin-coated coverslips. Quantification of the numbers of DC1-OT-I conjugates generated and percentage of DC1 conjugating (n = 10 planes per group), surface of interaction at the IS (actin maximum angle), and actin thickness at the IS (n = 20-32 DC1-OT-I conjugates per group). One-way ANOVA followed by Tukey's post-test. **C.** Average actin distribution in control and tumor-conditioned lung DC1 (n = 10 cells per group). **D.** Representative confocal planes of control, early (d8) and late (d28) DC1 (scale bar, 4 µm). Quantification of the cortical actin thickness (n = 15 cells per group). One-way ANOVA followed by Tukey's post-test. **E.** Isolated lung DC1 were loaded *ex vivo* with different concentrations of SIINFEKL and mixed with CTV-labelled OT-I cells (1:10 DC1:T cell ratio). Representative heatmaps showing the percentage of OT-I cells in each proliferation cycle after 72 hs of co-incubation at 0 (no peptide), 0.3 and 1 nM SIINFEKL. Data represent mean ± SEM. **p<0.01 ****p<0.0001.

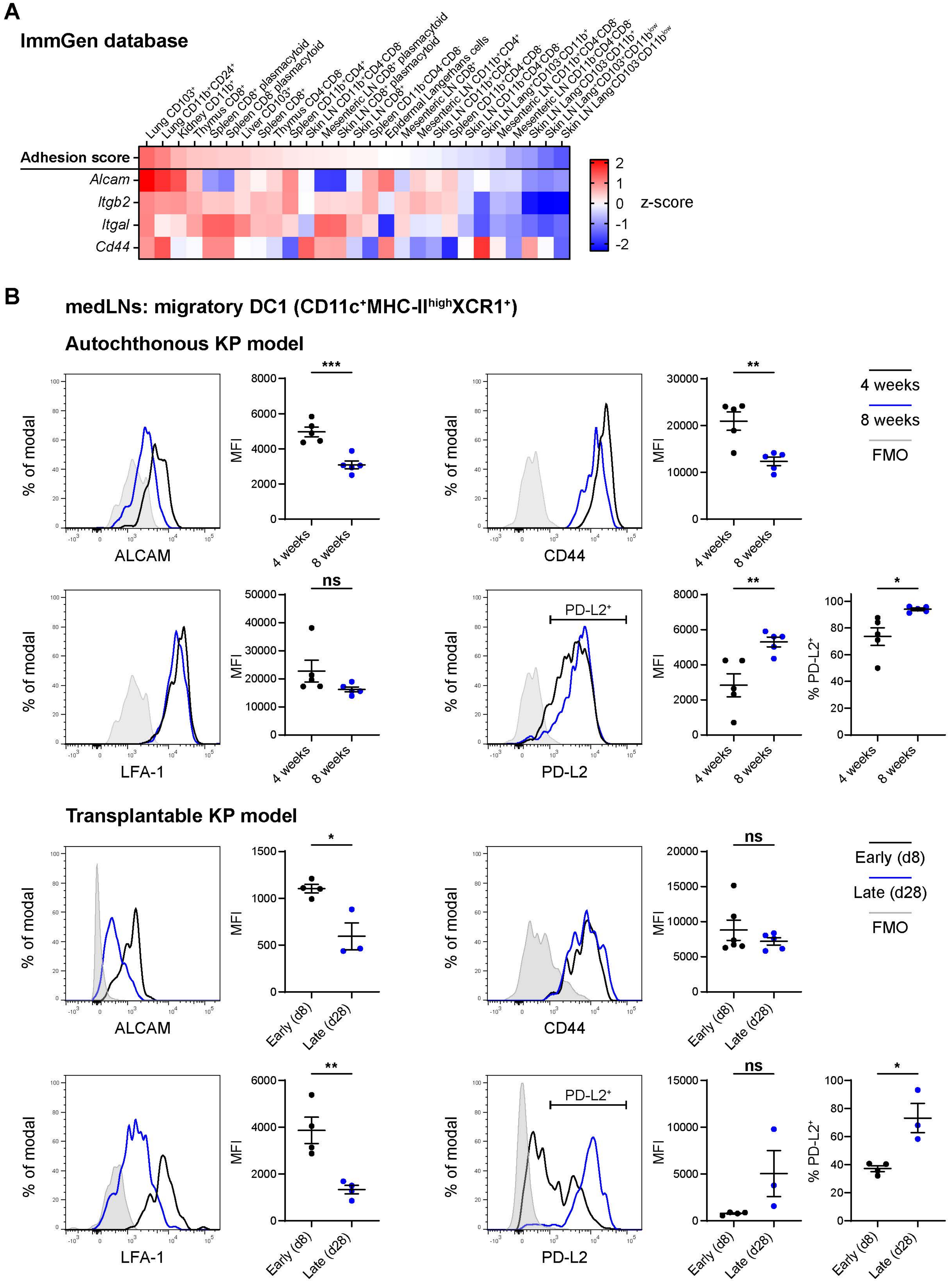
A
ImmGen database
Lung CD103+
Lung CD11b+CD24+
Kidney CD11b+
Thymus CD8+
Spleen CD8+ plasmacytoid
Spleen CD8- plasmacytoid
Liver CD103+
Spleen CD8+
Thymus CD4-CD8-
Spleen CD11b+CD4+
Skin LN CD11b+CD4-CD8-
Mesenteric LN CD8+ plasmacytoid
Skin LN CD8+ plasmacytoid
Skin LN CD8+
Spleen CD11b-CD4-CD8-
Epidermal Langerhans cells
Mesenteric LN CD8+
Mesenteric LN CD11b+CD4+
Skin LN CD11b+CD4+
Spleen CD11b+CD4-CD8-
Skin LN CD11b-CD4-CD8-
Skin LN Lang+CD103-CD11b+
Mesenteric LN CD11b+CD4-CD8-
Mesenteric LN CD11b-CD4-CD8-
Skin LN Lang-CD103-CD11b+
Skin LN Lang+CD103+CD11blow
Skin LN Lang-CD103-CD11blow
Adhesion score
Alcam
Itgb2
Itgal
Cd44
z-score
B
medLNs: migratory DC1 (CD11c+MHC-IIhighXCR1+)
Autochthonous KP model
% of modal
ALCAM
MFI
4 weeks
8 weeks
***
CD44
**
4 weeks
8 weeks
FMO
LFA-1
ns
PD-L2+
PD-L2
**
*
% PD-L2+
Transplantable KP model
*
Early (d8)
Late (d28)
ns
FMO
**
ns
*

Figure S2

**Figure S2. Downregulation of ALCAM and LFA-1 in advanced KP tumors is also mirrored by migratory DC1 in tumor-draining lymph nodes. A.** Heatmap showing the expression of *Alcam*, LFA-1 (*Itgab2, Itgal*), and *Cd44* across steady-state DCs in diverse tissues and conditions, taken from the ImmGen Database. DCs were ordered based on an Adhesion score calculated as the mean z-score between these genes. **B.** Mediastinal lymph nodes (medLNs) from tumor-bearing mice were harvested and processed for flow cytometry. In the autochthonous model, medLNs were harvested and analyzed after 4- (black line) or 8-weeks (blue line) post Ad-Cre administration (n = 4 mice per group, one representative out of two independent experiments). In the transplantable model, medLNs were analyzed at day 8 (black line, n = 4-6 mice) or 28 (blue line, n = 3-5 mice) after intravenous KP tumors inoculation (one representative out of three independent experiments). Fluorescence minus one (FMO) control staining for each marker is shown in grey. Quantification of the Median Fluorescence Intensity (MFI) or percentage of positive cells in lung DC1 for the indicated markers is presented. Data represent mean ± SEM. Unpaired Student t-test. $^*p<0.05$ $^{**}p<0.01$.
